## Supplementary material for "Highly-multiplexed serology for non-human mammals": S1 File

### S1 File. Pan-coronavirus PepSeq Library

Our pan-coronavirus PepSeq assay (PCV) was designed to broadly cover potential linear epitopes from all sequenced coronaviruses. On March 2, 2020, we downloaded all viral protein sequences from the UniProt Knowledgebase ("uniprot_sprot_viruses.dat" and "uniprot_trembl_viruses.dat" from https://ftp.uniprot.org/pub/databases/uniprot/current_release/knowledgebase/taxonomic_divisions/), and extracted 37,022 sequences annotated with six target taxonomy IDs: NCBI:txid693996 (*Alphacoronavirus* genus), NCBI:txid694002 (*Betacoronavirus* genus), NCBI:txid694013 (*Gammacoronavirus* genus), NCBI:txid1159901 (*Deltacoronavirus* genus), NCBI:txid693995 (*Coronavirinae* subfamily), and NCBI:txid1986197 (unclassified *Coronaviridae* family). We removed all sequences <30 amino acids in length and collapsed identical sequences to a single representative using a custom python script (https://github.com/LadnerLab/Library-Design/tree/master/one_hundred_reps/python). Prior to clustering, we attempted to assign proteins linked to ambiguous taxonomic IDs (NCBI:txid693995 and NCBI:txid1986197) to one of the four genera in the *Coronaviridae* family. These assignments were made based on kmer similarity to proteins already linked to these genus-level IDs using a custom python script (https://github.com/LadnerLab/PepSIRF/blob/master/extensions/checkTaxonomy.py). Specifically, to assign one of these ambiguous sequences to a genus, we required it to share at least 20% of its 4mers with a genus-linked protein, and using this criteria we were able to putatively assign >98% (1279/1300) of the ambiguous sequences.

We then used Usearch v.10.0.240 [1] to cluster similar proteins. Usearch was run separately for each genus and for those that were not able to be assigned to a genus. The cluster_fast command was used with an identity threshold of 0.5. For clusters containing >15 sequences, we designed 30mer peptides using our epitope-centric set cover design algorithm (https://github.com/LadnerLab/Library-Design/tree/master/setCover/c), with settings that assured inclusion of 90-93% of all unique, potential 9mer epitopes. For clusters containing <15 sequences, we aligned the sequences from each cluster using Muscle v3.8.31 [2] and designed 30mer peptides using a sliding window approach, with a step size of 22, which ensures inclusion of all potential 9mer epitopes (https://github.com/LadnerLab/Library-Design/tree/master/slidingWindow/python). In addition, because there were few SARS-CoV-2 protein sequences present in Uniprot at the time that we downloaded our target sequences, we supplemented our PCV design with peptides designed using our set cover algorithm and targeting the 2309 SARS-CoV-2 genomes used for our SCV2 design described in Ladner et al. [3]. We considered the full SARS-CoV-2 proteome for this design and used settings that ensured inclusion of 100% of the unique, potential 12mer epitopes. Our final PCV design included 97,699 peptides generated from coronavirus proteins downloaded from Uniprot, 1796 peptides generated from SARS-CoV-2 proteins, and 476 control peptides designed from a variety of viruses (not coronaviruses) and known to be commonly recognized by

IgG antibodies in human sera [3]. Each peptide was chosen to be 30 amino acids in length and each was represented by a single nucleotide encoding.


1. Edgar RC. Search and clustering orders of magnitude faster than BLAST. Bioinformatics. 2010;26: 2460–2461.

2. Edgar RC. MUSCLE: multiple sequence alignment with high accuracy and high throughput. Nucleic Acids Research. 2004. pp. 1792–1797. doi:10.1093/nar/gkh340

3. Ladner JT, Henson SN, Boyle AS, Engelbrektson AL, Fink ZW, Rahee F, et al. Epitope-resolved profiling of the SARS-CoV-2 antibody response identifies cross-reactivity with endemic human coronaviruses. Cell Rep Med. 2021;2: 100189.

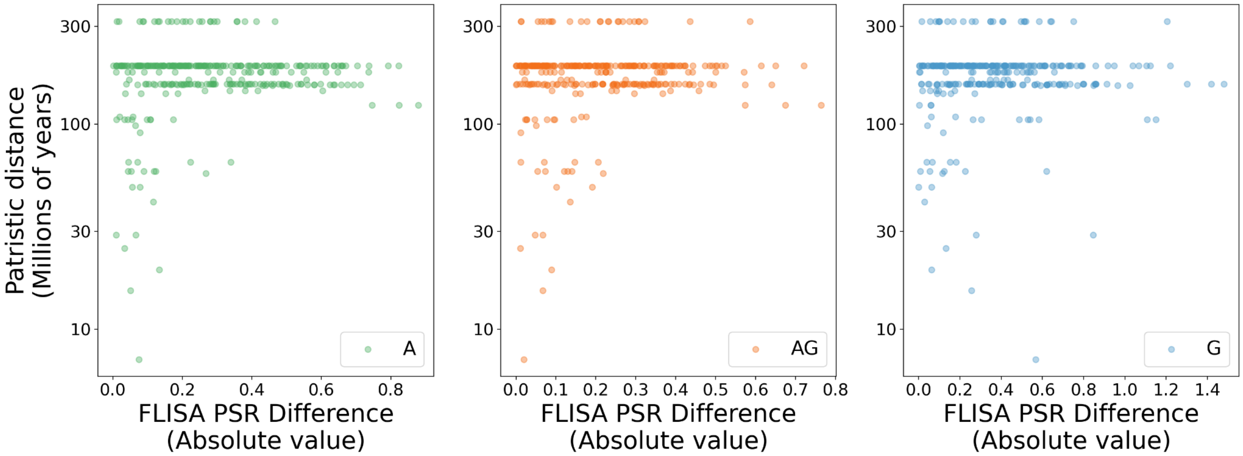


**S1 Figure. Species with close phylogenetic relationships tend to exhibit similar estimated binding affinities to the IgG-binding proteins tested in this study.** Scatterplots comparing pairwise differences in FLISA positive slope ratio (PSR) values measured in this study (x-axis, absolute values shown) and the sum of the length of branches separating species in a time-scaled phylogeny (y-axis, calculated from tree shown in Figure 2). Each point represents a pairwise comparison between species, with a total of 26 species included (325 pairwise comparisons). Each panel represents a different IgG-binding protein: protein A (left, green), protein AG (middle, orange), and protein G (right, blue).

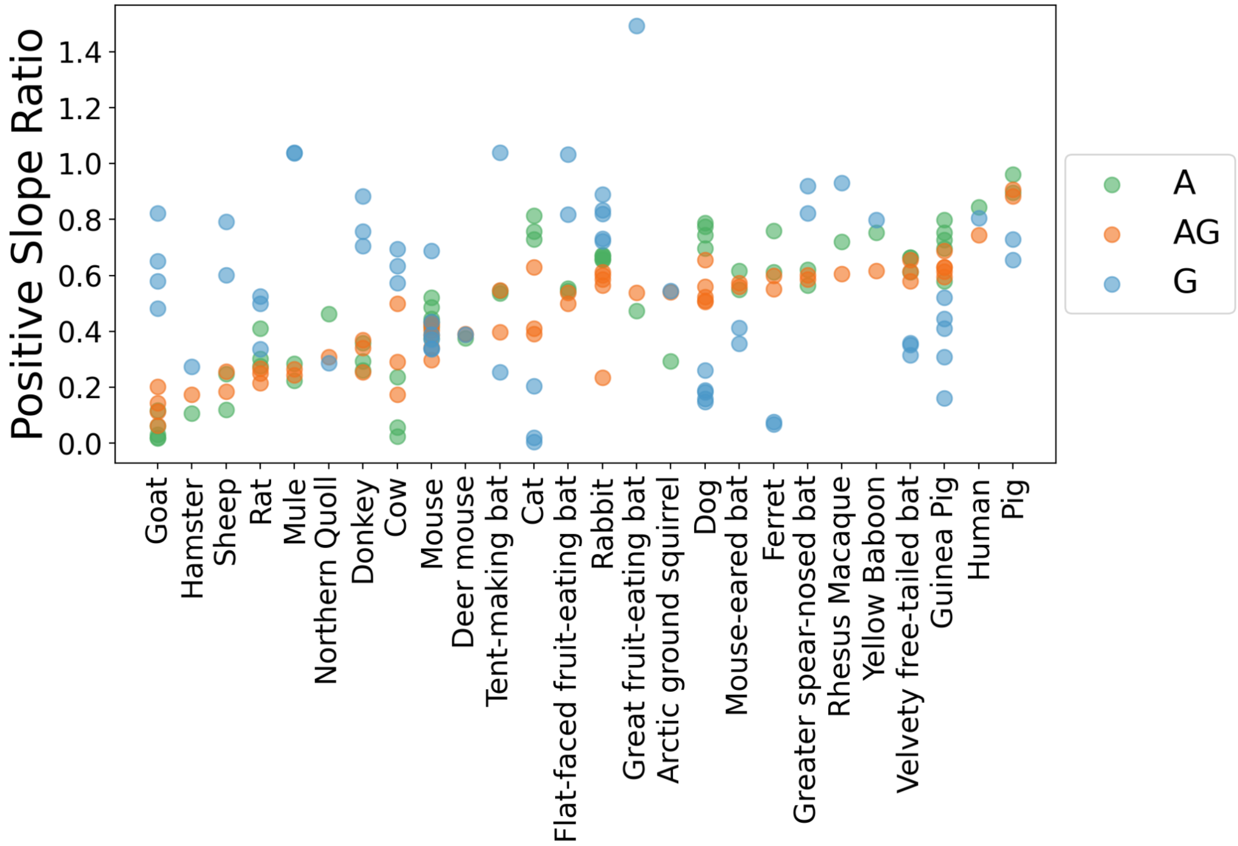


**S2 Figure. Relative binding affinities measured for protein AG (orange) were often intermediate between affinities measured for protein A (green) and protein G (blue).** FLISA positive slope ratio values measured for the 26 mammal species characterized in this study. Species are sorted along the x-axis according to the average positive slope ratio for protein AG (ascending order). Each point represents an individual sample and IgG-binding protein combination (64 data points per IgG-binding protein). For scientific names of each species, see S1 Table.

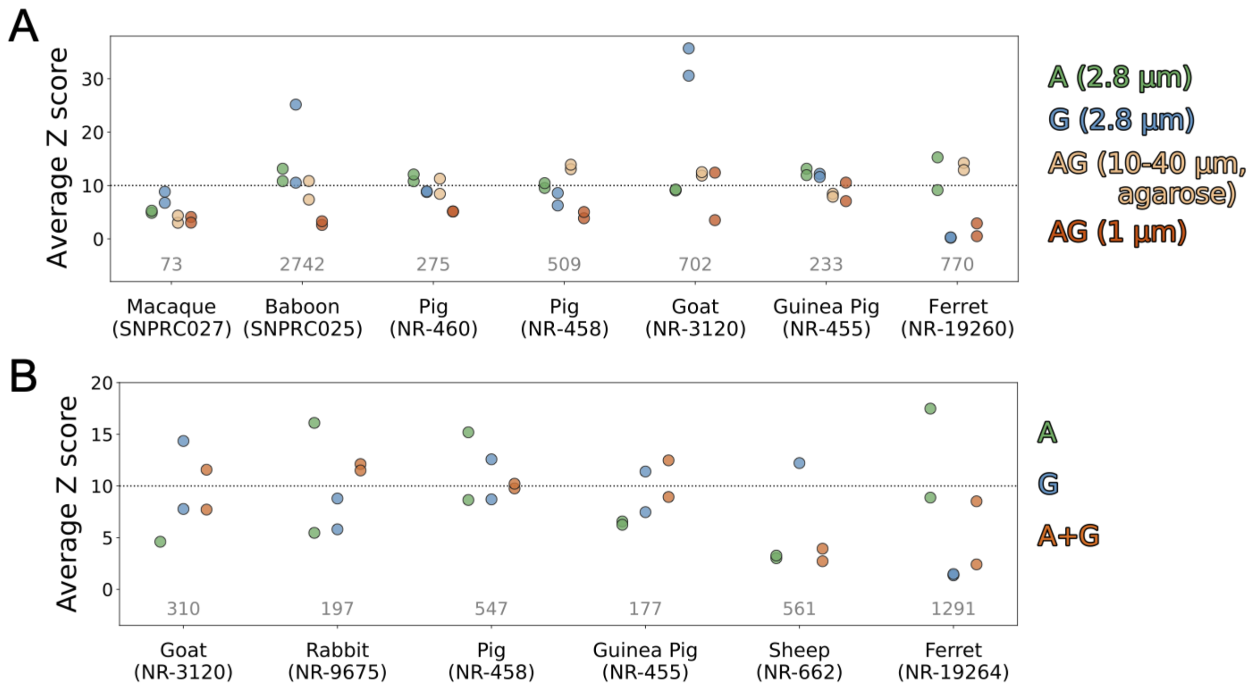


**S3 Figure. Average Z scores from PepSeq assays using protein A and protein G IgG-binding domains as capture proteins, both in isolation and in combination.** Each point represents results from a single HV1 PepSeq assay. Most sample/capture protein combinations are represented by two replicates, but some are only represented by one because one of the replicates failed quality control thresholds. Differences between replicates represent technical variation in enrichment signal. (A) Enrichment signal comparison between protein A (green), protein G (blue) and recombinant protein AG (orange), which is a recombinant protein that consists of fused IgG-binding domains from both protein A and protein G. Two different bead types were tested for protein AG: 1 µm magnetic beads (Pierce #88802, dark orange) and 10-40 µm agarose magnetic beads (Pierce #78609, light orange). (B) Enrichment signal comparison between protein A (green), protein G (blue) and a combination of protein A and protein G bearing magnetic beads (orange). All of the assays from a given panel were run on a single 96-well plate. Average Z scores for a given sample/panel were calculated considering all of the peptides, across the tested capture proteins, with a Z score ≥10 in ≥1 replicate. The number of considered peptides for each sample is shown in gray.

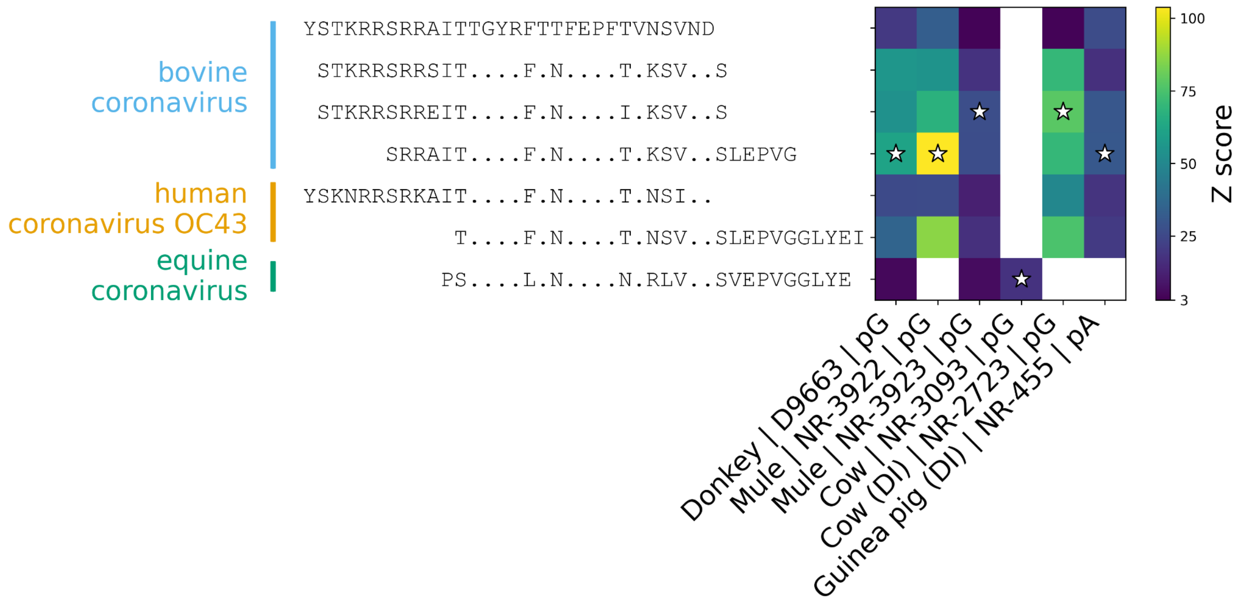


**S4 Figure. PepSeq enrichment Z scores for diverse peptides covering a single epitope region of the betacoronavirus 1 Spike protein.** Our HV1 PepSeq assay included seven peptides derived from viruses of the betacoronavirus 1 species and overlapping a commonly reactive antibody epitope at the S1/S2 cleavage site. This included 4, 2 and 1 peptide(s) derived from bovine coronavirus, human coronavirus OC43 and equine coronavirus, respectively. The heat map shown here illustrates relative enrichment Z scores across these seven peptides for all six samples with ≥1 enriched peptide at this epitope (see Figure 5B). For each sample, the white star indicates the peptide with the highest Z score. To the left of the heat map is a multiple sequence alignment of these seven peptide sequences, where periods (".") indicate sites that are identical across all seven peptides. X-axis labels indicate the species from which the sample was obtained, the sample ID, and the capture protein used in the PepSeq assay (pA = protein A, pG = protein G).

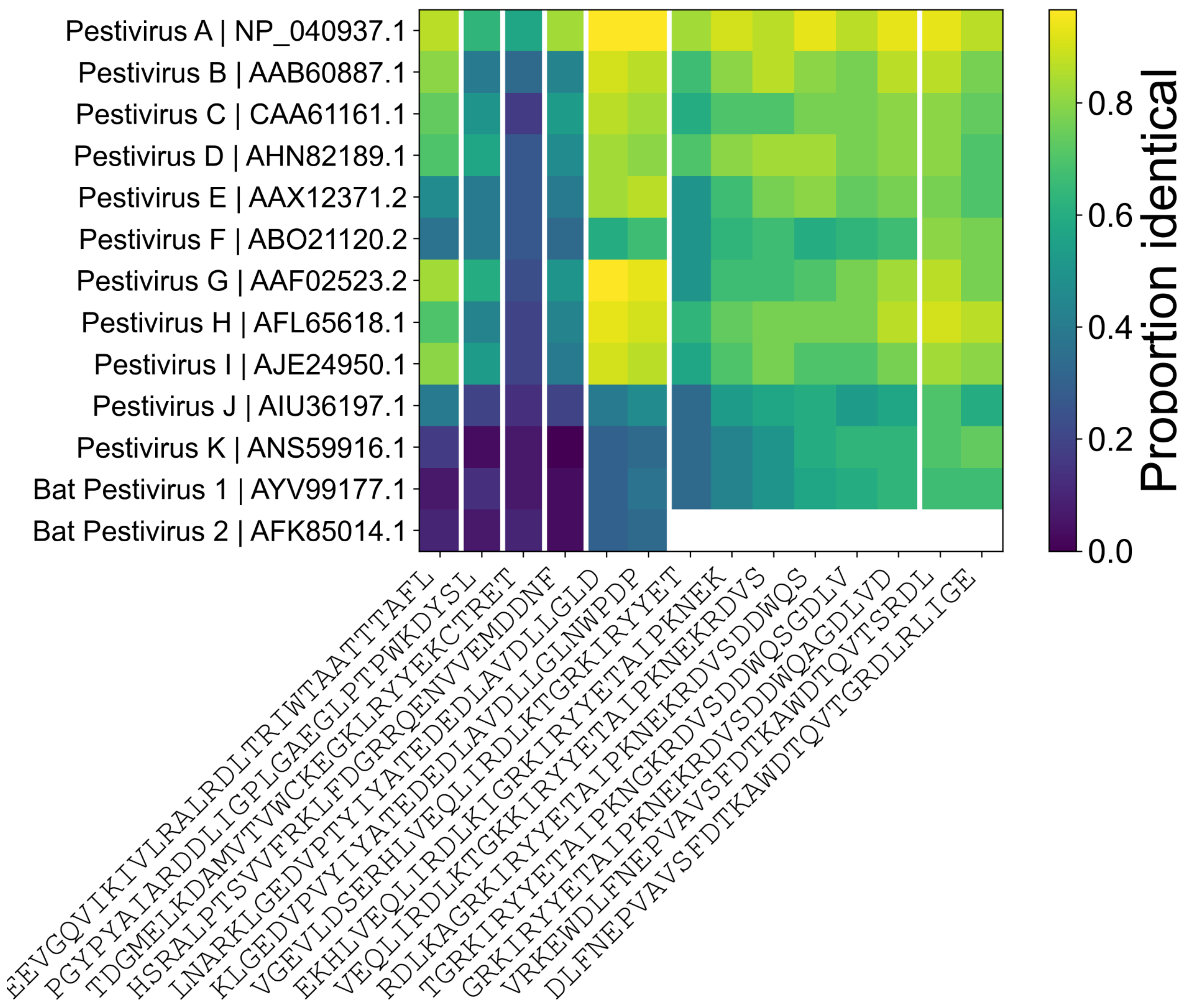


**S5 Figure. Sequence conservation (amino acid level) across 14 peptides (x-axis) and 11 species and two unclassified viruses within the *Pestivirus* genus (y-axis).** In our HV1 library PepSeq assays of sera collected from bats in Necoclí-Colombia, we detected a total of 14 enriched pestivirus A peptides across eight reactive samples (see Figure 5C). Pestivirus A is the only species within the *Pestivirus* genus that is included in our HV1 assay. Therefore, we can not directly assess whether this antibody reactivity is likely to represent true exposure to this species or whether these could reflect cross-reactive antibodies stimulated by infection with a different pestivirus species. To assess the level of sequence conservation at these antibody epitopes, we aligned representative polyprotein sequences for all 11 named pestivirus species (A-K) and two unclassified viruses sequenced from bats (species names and GenBank accession #s provided in y-axis labels). Then, for each enriched peptide, we calculated the proportion of identical residues within each of these representative sequences. For three of the epitopes, we observed multiple, enriched peptides. Vertical white bars separate each epitope region. The available sequence for bat pestivirus 2 (GenBank: AFK85014.1) is incomplete, which is why there are no values associated with the last two epitope regions (8 peptides) for this virus.
