## Supplementary figures and images for "Highly-multiplexed serology for non-human mammals"

### S1 Figure

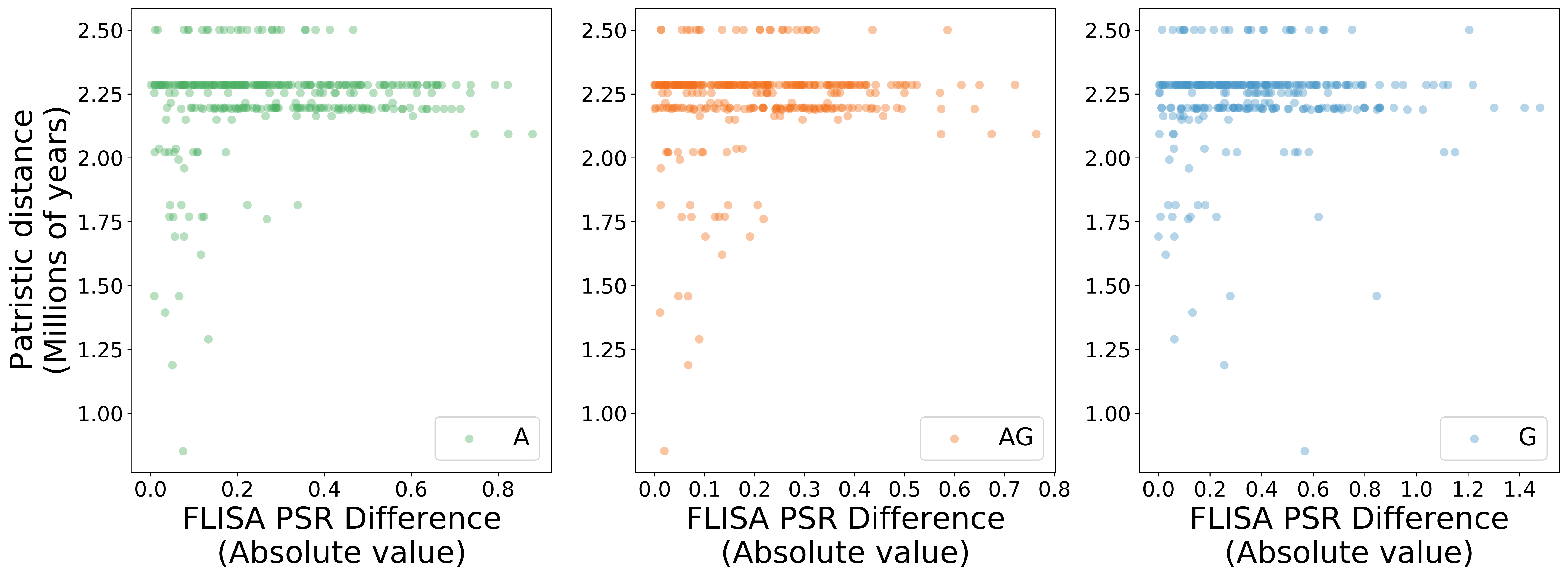

### S2 Figure

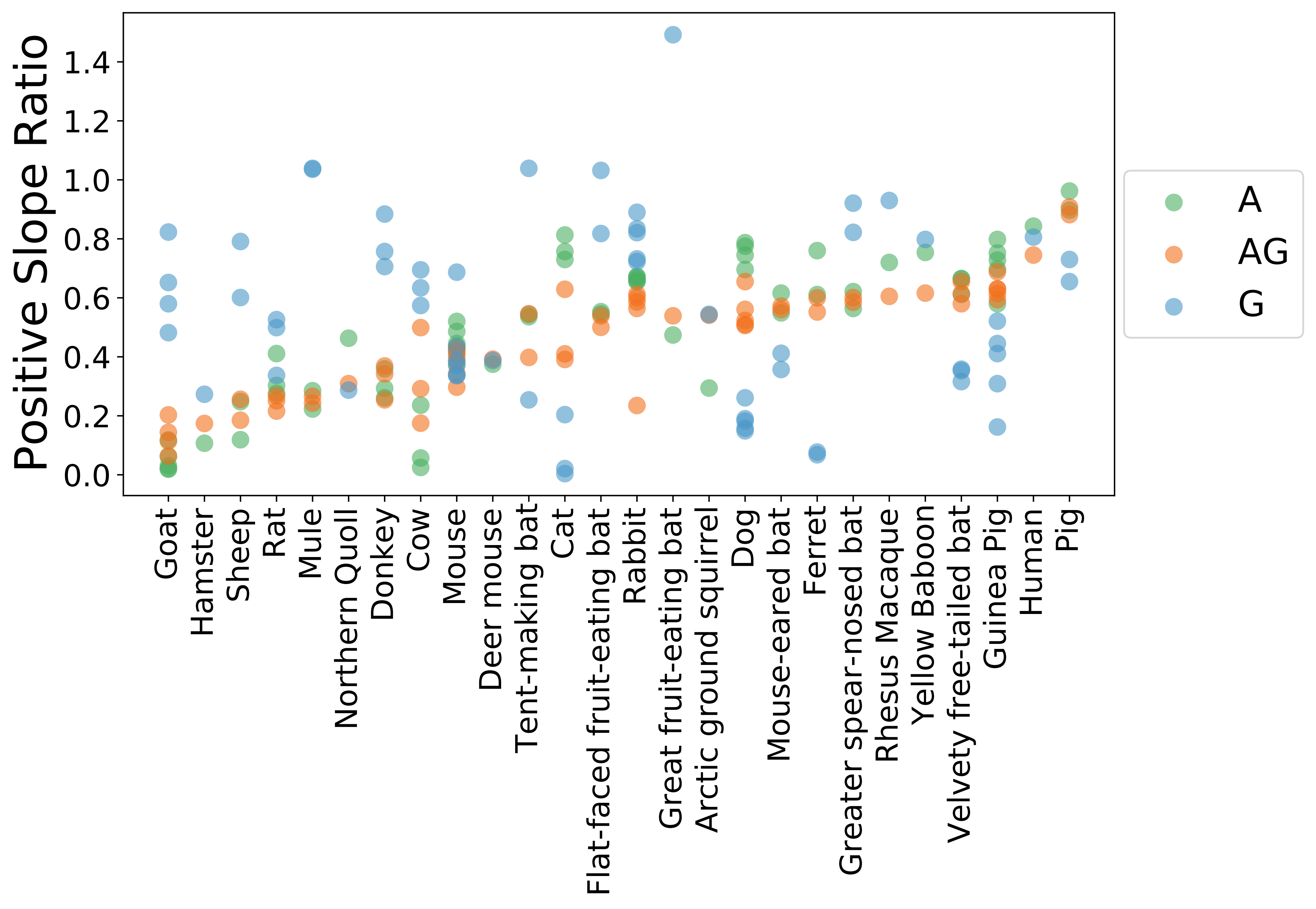

### S3 Figure

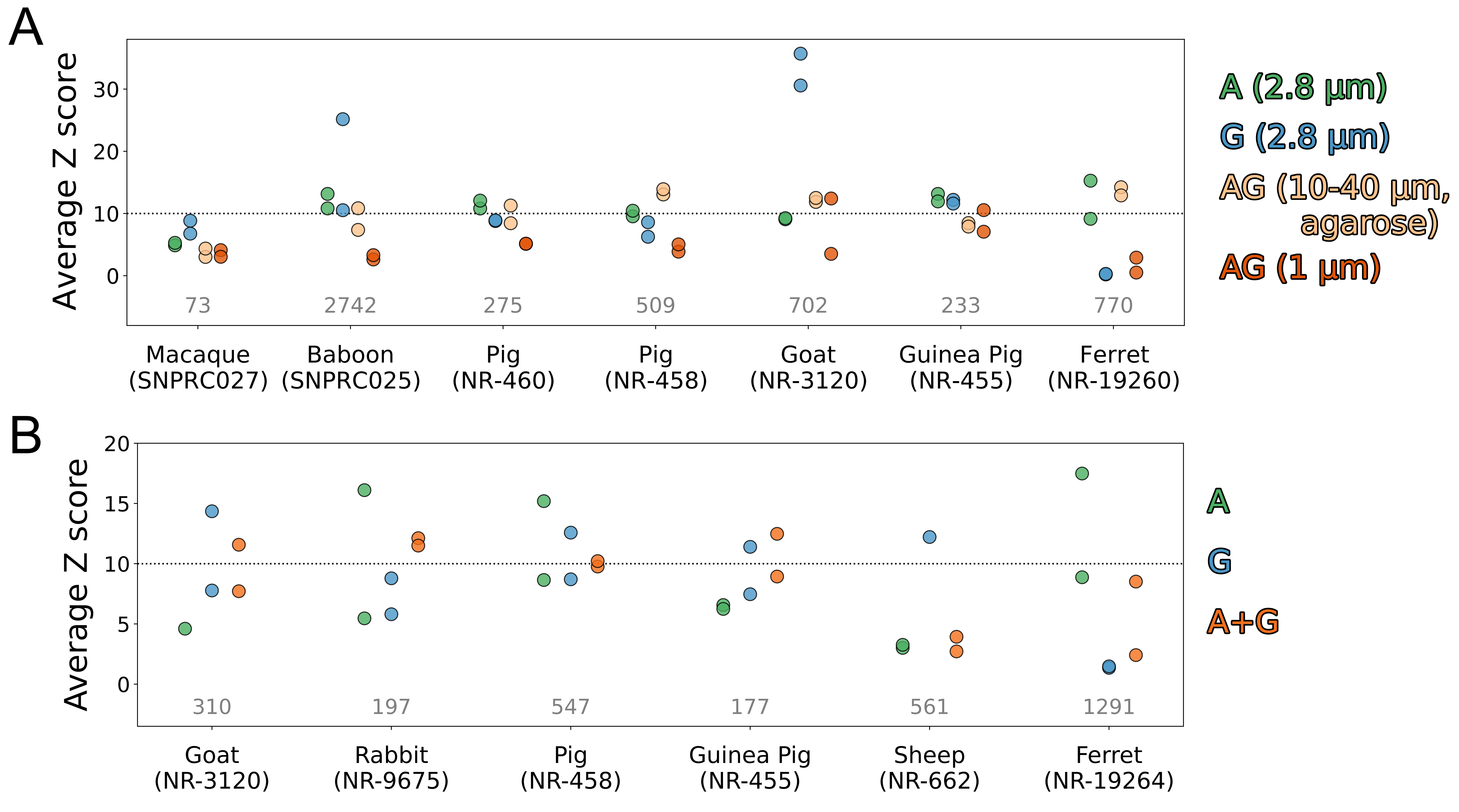

### S4 Figure

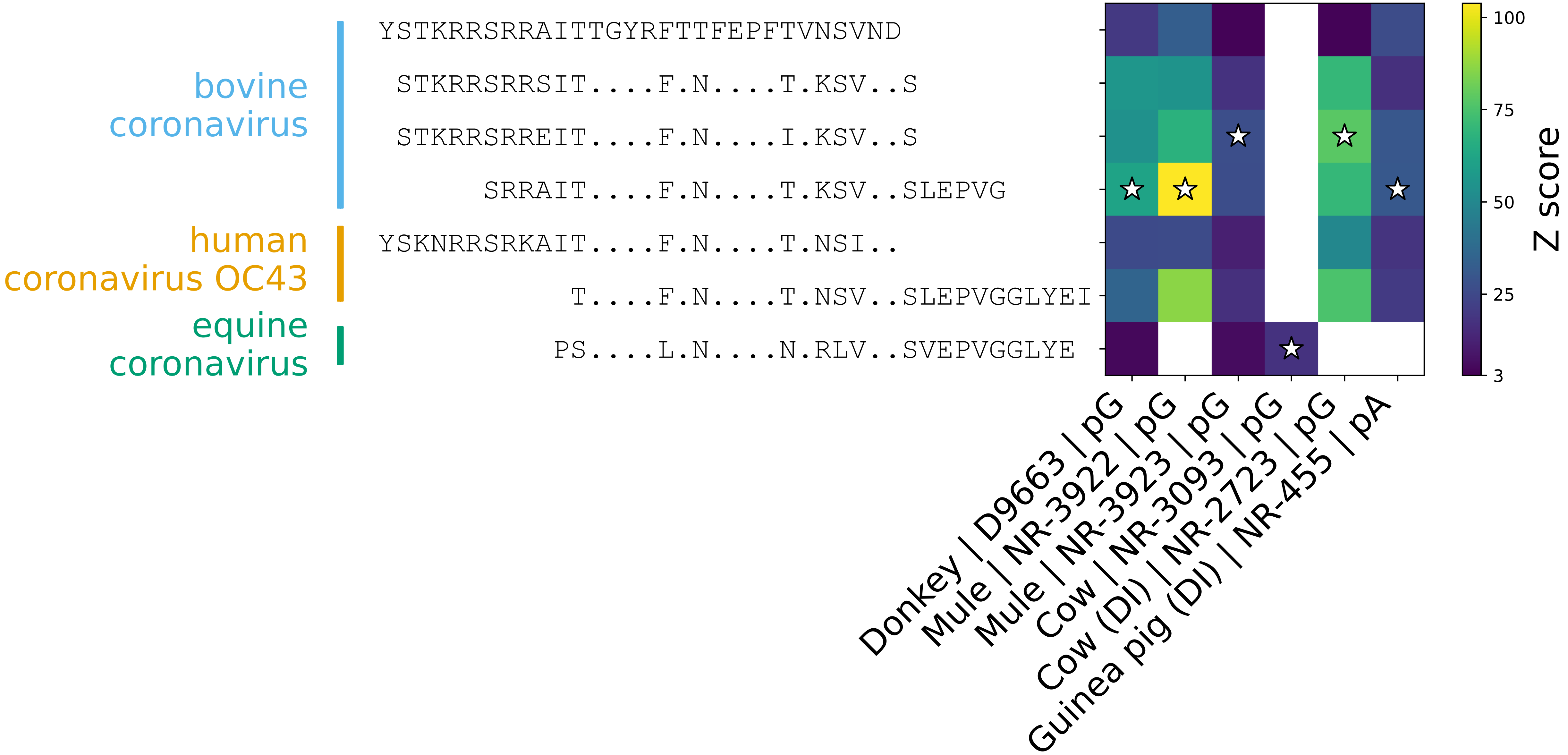

### S5 Figure

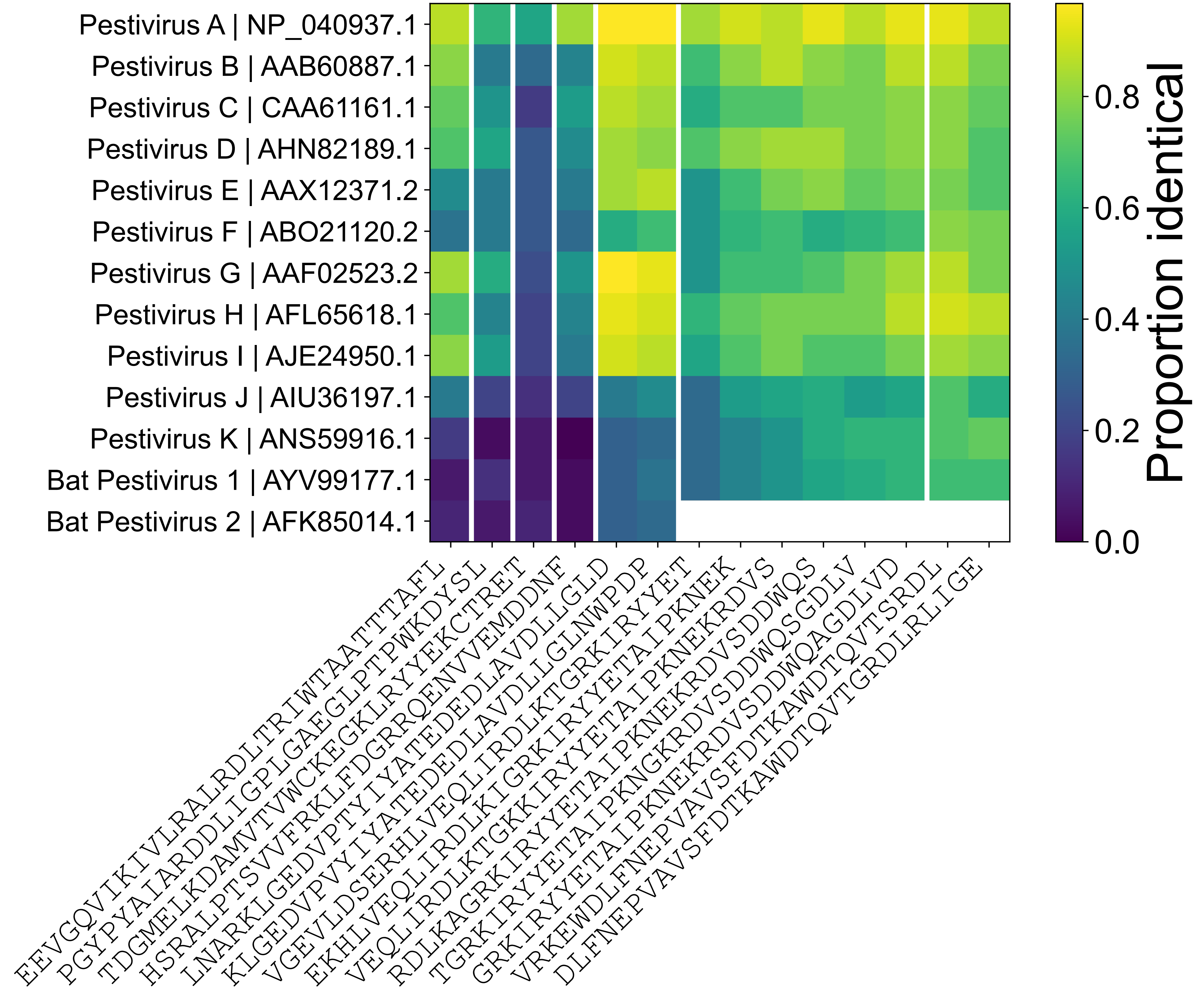
